# Evidence implicating sequential commitment of the founder lineages in the human blastocyst by order of hypoblast gene activation

**DOI:** 10.1101/2022.12.08.519626

**Authors:** Elena Corujo-Simon, Arthur H. Radley, Jennifer Nichols

## Abstract

Successful human pregnancy depends upon rapid establishment of three founder lineages: trophectoderm, epiblast and hypoblast, which together form the blastocyst. Each plays an essential role in preparing the embryo for implantation and subsequent development. Several models are proposed to define the lineage segregation. The first suggests that all specify simultaneously; the second favours differentiation of trophectoderm before separation of epiblast and hypoblast, either via differentiation of hypoblast from established epiblast, or production of both tissues from the inner cell mass precursor. To begin to resolve this discrepancy and thereby understand the sequential process for production of viable human embryos, we investigated the expression order of genes associated with emergence of hypoblast. Based upon published data and immunofluorescence analysis for candidate genes, we present a basic blueprint for human hypoblast differentiation, lending support to the proposed model of sequential segregation of the founder lineages of the human blastocyst. The first characterised marker, specific initially to the early inner cell mass, and subsequently identifying presumptive hypoblast is PDGFRA, followed by SOX17, FOXA2 and GATA4 in sequence as the hypoblast becomes committed.

**Summary Statement:** Optimal segregation of human blastocyst founder lineages is essential to establish healthy human pregnancies. Mapping activation of hypoblast marker genes over time helps understand how the yolk sac is regulated.

## Introduction

At the time of implantation in the uterus, around 8 days after fertilisation (D8), the human embryo comprises three lineages: trophectoderm (TE), epiblast (epi) and hypoblast (hypo). TE is responsible for connecting to the uterus and is the precursor of the placenta. Epi gives rise to the embryo proper and hypo will form the primary yolk sac (Hertig et al., 1956). Hypo also plays an essential role in patterning the epi to establish the foetus (Mole et al., 2021). The mechanisms by which the three lineages segregate have been well described for the mouse embryo, but emerging data from human embryos implies that the developmental processes may not be completely conserved between the species. Successful pregnancy rate following assisted conception programmes is disappointingly low (around 25% of transferred embryos, according to HFEA metrics). Understanding how the lineages segregate and quantifying the proportions of each required for mature blastocyst morphology constitute prerequisites for optimising culture regimes to improve live birth rates following IVF treatment. So far, several models have been proposed to explain acquisition of the three pre-implantation lineages in the human embryo. The one-step model suggests that TE, epi and hypo cell fates are acquired synchronously at around D5, based on analysis of single cell RNA sequencing (scRNAseq) data (Petropoulos et al., 2016). In contrast, the two-step model proposes that, in concordance with the mouse embryo, TE and inner cell mass (ICM) lineages begin to segregate at the morula stage, with the hypo and epi emerging subsequently from the ICM following cavitation of the blastocyst (Blakeley et al., 2015; Niakan and Eggan, 2013; Roode et al., 2012). Other scRNAseq analyses and immunofluorescence (IF) for pan markers of TE versus ICM studies support position initiation of the TE programme at the morula stage (Gerri et al., 2020; Meistermann et al., 2021; Stirparo et al., 2018; Zhu and Zernicka-Goetz, 2020), prior to appearance of epi and hypo as the blastocyst expands. These reports indicated that a combination of polarity acquisition and activation of phospholipase C and Hippo signalling leads outer morula cells to commence TE differentiation with activation of GATA3 and nuclear localisation of YAP. Subsequently, it was suggested that hypo originates from the epi (Meistermann et al., 2021). However, a more recent study, using an entropy-based approach for feature selection for published scRNA-seq data sets identified an early ICM population from which both epi and hypo arise (Radley et al., 2022). The existence of this common ICM precursor had also been implicated previously (Stirparo et al., 2018).

The process of epi and hypo specification in human embryos has not yet been described in detail, although it is known that hypo is not induced simply as a result of FGF/ERK signalling from the epi, as it is in the mouse (Kuijk et al., 2012; Nichols et al., 2009; Roode et al., 2012). Divergence in the order of gene expression during this segregation in human compared to mouse was first investigated using IF (Niakan and Eggan, 2013) and subsequently expanded with scRNAseq, enabling identification of 164 genes differentially expressed between the two lineages (Yan et al., 2013). A larger dataset identified *LINC00261* as the most highly expressed hypo gene; *PDGFRA, FGFR2, LAMA4, HNF1B, COL4A1, GATA4, FN1, FRZB, AMOTL1*, and *DPPA4* were also highlighted (Petropoulos et al., 2016). Hypo markers were further separated as ‘early’ for those expressed before the hypo is fully specified (*GATA6, LRP2* and *ANXA3)* and ‘late’ for genes appearing only in the mature lineage (*APOA1, COL4A1, GDF6, RSPO3* and *FST)*. OTX2 was also discovered to mark early human hypo, providing another example of divergence in lineage identity from that of mouse (Boroviak et al., 2018).

In this study, we sought to specify the order of appearance of hypo-associated gene products and thereby further delineate blastocyst staging and enrich embryo quality control. We compare initiation and downregulation of candidate hypo genes and proteins during blastocyst expansion alongside known epi markers to resolve the sequence of cell fate specification in the human embryo and thereby determine whether hypo cells arise simultaneously with the other two founder lineages, via conversion of epi-specified cells, or from the pluripotent ICM following its segregation from TE.

## Results and Discussion

### The hypoblast lineage is acquired progressively, concurrent with intensification of SOX17 in ICM cells during blastocyst development

To study acquisition of hypo and epi fates in the human embryo, we thawed D5 blastocysts and fixed them immediately after recovery, or following culture to ‘early’ D6, ‘late’ D6 or D7 stages. Although still present in the TE at early stages, OCT4 was used to track the ICM/epi lineage, while SOX17 identified hypo (Fig. 1A) (Niakan and Eggan, 2013). Embryos were staged using a combination of number of days’ culture post-fertilisation and morphological landmarks, including thickness of zona pellucida, size of blastocoel and total cell number (Fig. S1). At D5, the embryo comprises mostly OCT4+ cells; the few SOX17+ cells also display high levels of OCT4 (Fig. 1A-D; Fig. S2). Notably, OCT4 is still present in TE at a similar level to that of the ICM at this stage. Consequently, the ICM/epi quantification also includes some TE cells (Fig. 1B-C; Fig. S2). At early D6, the majority of ICM cells co-expresses OCT4 and SOX17; detection of cells displaying high levels of OCT4 without SOX17 suggests emergence of epi precursors (Fig. 1A-D). At late D6, the ICM can be separated into three populations of cells: epi precursors expressing only OCT4; ‘undefined’ ICM cells displaying high levels of SOX17 as well as OCT4, and hypo precursors expressing SOX17, but no OCT4 (Fig. 1A-D). At D7, the undefined population has resolved into epi and hypo cells, distinguishable by expression of either OCT4 or SOX17. Coincidentally, by this stage the hypo is located near the blastocoel (Fig. 1A). This sequence of events suggests that, in a similar way to the mouse embryo, nascent human ICMs (D5) possess some OCT4+ SOX17-cells, whereas by early D6, cells co-express markers for epi and hypo, indicating presence of dual progenitor cells within the ICM population with the potential to form either epi or hypo. The number of OCT4+SOX17+ cells per embryo increases at early D6 as SOX17 expression begins in ICM cells (Fig. 1E, Fig. S2). The number of double positive cells reduces at late D6 as OCT4 intensity diminishes in the hypo precursors, leading to complete segregation of hypo from epi by D7 (Fig. 1B-C, E-F, Fig. S2). This analysis was repeated in three more datasets showing the same trend: starting from low or no SOX17 expression in the ICM, to OCT4+SOX17+ co-expression and finally segregation of ICM cells into exclusively OCT4 (epi) or SOX17 (hypo) lineages (Fig. S2).

**Figure 1.**
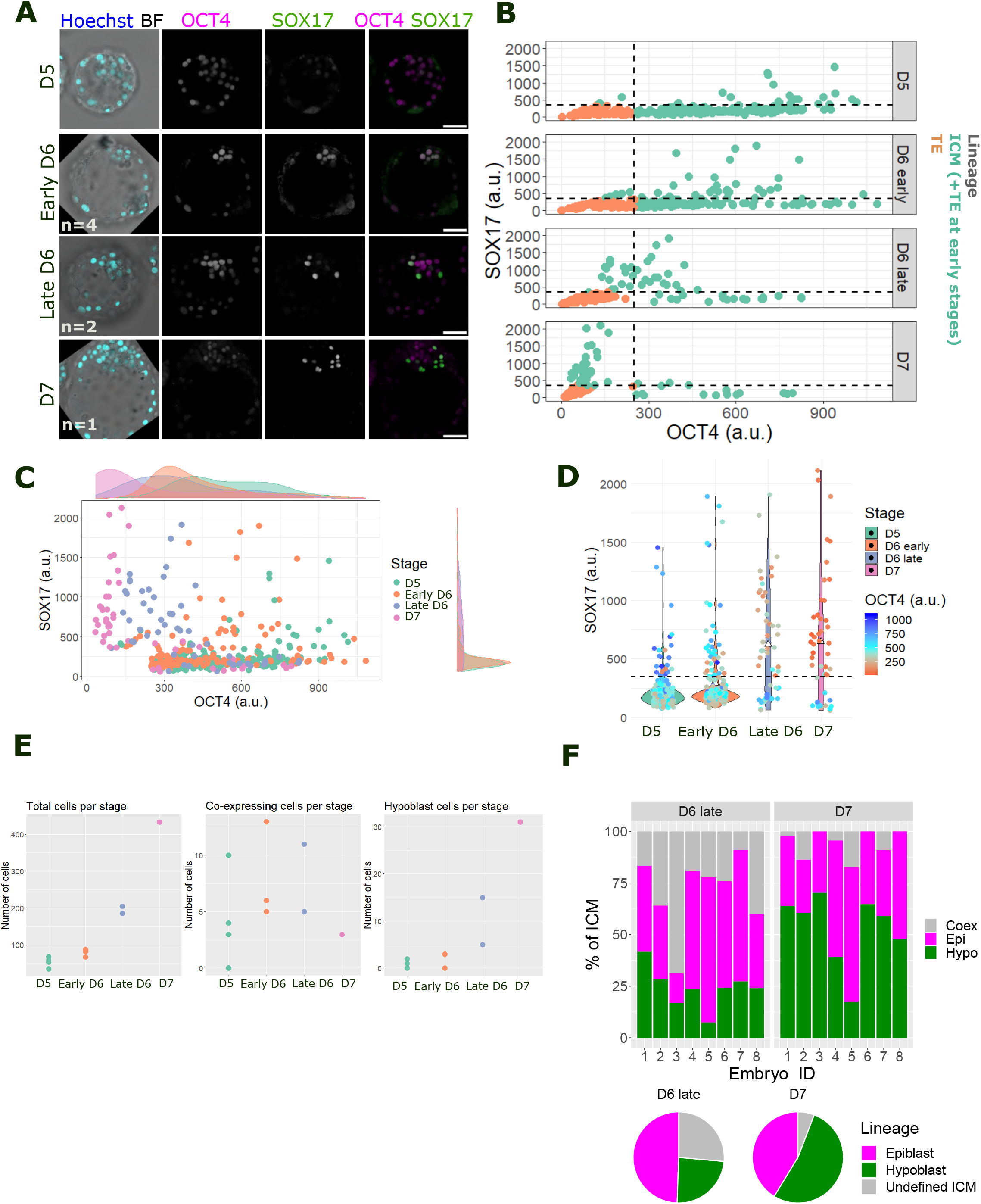
Tracing appearance of hypoblast in the human blastocyst from D5 to D7 based on co-staining of SOX17 and OCT4. **A**. Representative confocal images of human blastocysts at day (D)5, early D6, late D6 and D7 immunostained for the epiblast marker OCT4 (magenta) and the hypoblast marker SOX17 (green). Images are a stack combination of 9 consecutive 1 µm single z images. “n” refers to number of embryos analysed per stage. Scale bar = 50 µm. **B**. Scatter plots quantifying the nuclear intensity level of OCT4 (x-axis) and SOX17 (y-axis) across blastocyst stages in the ICM (light green) and TE (orange). At early stages (D5 and early D6), light green also comprises TE cells expressing OCT4 at the same level as ICM. Each dot corresponds to one cell. Dashed lines represent intensity level/expression threshold calculated at D7 when TE and ICM (epiblast and hypoblast) are completely segregated. **C**. Scatter plot combining nuclear intensity data from ICM cells at different stages. Marginal density plots showing the distribution of data into distinct discrete populations or showing heterogeneous levels. **D**. Violin plot comparing nuclear fluorescence intensity level of SOX17 across stages and its co-expression nuclear OCT4 per cell. The dashed line represents the expression threshold for SOX17 calculated at D7. **E**. Swarm plot for the absolute number of total cells (used from embryo staging), co-expressing and hypoblast cells per stage. **F**. Top panel: stacked bar plot quantifying the percentage of each lineage population in the ICM of individual embryos at late D6 and D7 stages. Lower panel: pie charts showing the average percentage of each lineage in each stage.

The proportion of epi and hypo (also called primitive endoderm) cells in mouse embryos at embryonic day E4.5 is largely consistent and maintained at 40:60% for all embryos (Saiz et al., 2020; Saiz et al., 2016). However, in D7 human embryos the proportion of cell lineages appears to vary from 20 to 60% for both hypo and epi (Fig. 1F). The percentage of putative epi cells in the ICM remains nearly constant from late D6 to D7 (49 to 41%), suggesting that it is the co-expressing cells at late D6 that will become hypo at D7 (Fig. 1F). Moreover, the small decrease in the percentage of epi cells implies they are fully specified and do not convert to hypo from late D6 to D7.

The progressive increase in hypo cells during blastocyst expansion and the scarcity of OCT4+SOX17+ co-expressing cells at D5 appears to support the hypothesis that hypo originates from epi cells in human embryos (Meistermann et al., 2021). In this case, exclusively OCT4+ epi cells at D5 will subsequently upregulate SOX17, generating the co-expressing population at D6 from which hypo differentiates at late D6 and D7. However, persistent OCT4 in the TE at this stage confounds the ability to identify epi cells definitively. Once co-expression is established at early D6, hypo cells apparently emerge from OCT4+SOX17+ cells rather than by direct conversion from Epi. It is possible that D5 OCT4+ cells express an alternative, earlier hypo marker distinct from SOX17, classifying them as early ICM cells (Radley et al., 2022). To test this hypothesis, we investigated alternative hypo markers expressed at D5 immediately after cavitation.

### Pseudotime analysis of blastocyst development unveils sequential activation of genes specifying the hypoblast lineage

In mouse blastocysts, hypo fate is acquired sequentially, originating from an ICM population co-expressing NANOG (epi) and GATA6 (hypo), which segregate into a ‘salt and pepper’ arrangement at around E3.5 (Plusa et al., 2008). Sequential activation of hypo markers was investigated using IF, revealing progressive expression of hypo transcription factors in the following order: GATA6 > SOX17 > GATA4 > SOX7 (Artus et al., 2011). This was subsequently confirmed by pseudotime analysis of scRNAseq data from early mouse embryos (Nowotschin et al., 2019). Hypo genes in human embryos were previously classified as either ‘early’ or ‘late’ using scRNAseq (Boroviak et al., 2018). Use of denoising and feature selection software (FFAVES and ESFW) expands the potential to detect small temporal differences in expression of individual genes (Radley et al., 2022). We therefore made use of this method to characterise the order of appearance of human hypo marker genes. The first step was to identify specific genes uniquely expressed in hypo and absent from TE and epi compartments. We began by selecting genes known to mark the mouse hypo population: *Pdgfra, Gata6, Sox17, Gata4* and *Sox7* (Artus et al., 2011; Chazaud et al., 2006; Plusa et al., 2008). To search for genes not yet known to be related to hypo, we leveraged a recently generated high resolution scRNAseq human pre-implantation embryo UMAP embedding (Radley et al., 2022) using published scRNAseq datasets (Fig. 2A-D, Fig. S3A,B). Genes were ranked based on level of enrichment in the hypo population according to the UMAP (Fig. 2D, Fig. S3C-F). Trophectoderm and epiblast genes were analysed as controls (Figure S3D-E). The order of appearance was obtained by fitting a logistic curve to smoothed gene expression values of the selected hypo genes along the hypoblast pseudotime and identifying the mid point of each logistic curve (Fig. 2C, Fig. S4). Based on these analyses, we suggest the following order: *PDGFRA > BMP2 > OTX2 > LGALS2 > VIL1 > SOX17 > FOXA2 > SALL1 > GATA4 > COL4A1 > RSPO3 > FRZB > IGF1 > FRLT3 > CPN1 > SYT13* (Fig. 2C-D).

**Figure 2.**
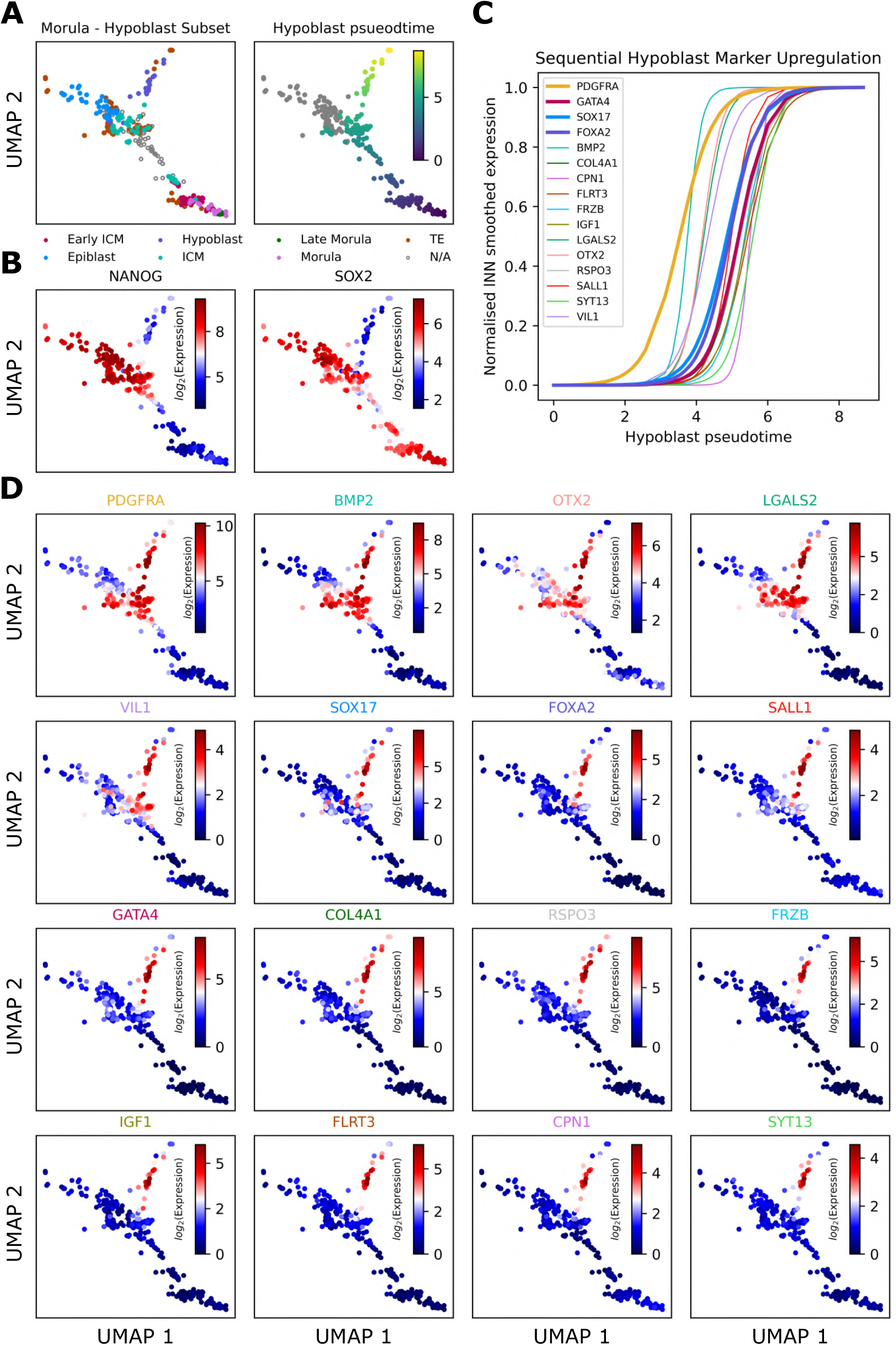
Pseudotime analysis of published scRNAseq from human embryos to establish the order of appearance of hypoblast markers during *ex vivo* development. **A**. The human pre-implantation embryo UMAP embedding described by Radley et al. 2022, sub-setted down to samples involved during the bifurcation of the morula to the epi/hypo populations. Left panel presents cell type labels defined by Stirparo et al. 2018. Right hand panel shows pseudotime along the hypo branch. **B**. NANOG and SOX2 expression distinguish hypo and epi populations. **C**. By fitting a logistic curve to the smoothed gene expression against the hypo pseudotime, we are able to delineate an ordering of hypo gene activation. **D**. Overlays of smoothed gene expression onto the UMAP confirms ordered upregulation specific to Hypo. Supplemental Fig 3 shows that these markers are exclusive to the hypo and not expressed in the TE.

The first divergence compared with mouse appeared with *GATA6 and SOX7. GATA6* appears to be a generic human preimplantation extraembryonic marker owing to its presence in both TE and hypo (Fig. S3D) (Boroviak et al., 2018; Roode et al., 2012). Surprisingly, *SOX7* is not expressed in human blastocysts, whereas *OTX2* marks hypo in human, but not mouse (Fig. S3F) (Boroviak et al., 2018); *FOXA2* appears in human hypo, but is absent in mouse at the mid blastocyst stage, according to scRNAseq. However, in late human blastocysts FOXA2 protein overlaps with SOX17, whereas it appears only in a subset of hypo cells at this stage in mouse (Blakeley et al., 2015). Thus, according to published data *PDGFRA, BMP2, OTX2* and *LGALS2* may be classified as early ICM genes; *VIL1, SOX17, FOXA2 and SALL1* as mid stage hypo genes and *GATA4* as a marker for mature hypo (Fig. 2, Fig. S3,4). To verify order of appearance of selected markers at the protein level we performed IF.

### PDGFRA is the first hypoblast marker to appear in the early ICM of human blastocysts

Validated antibodies for many of the novel hypo markers identified through the high resolution UMAP are not yet available. We therefore focused on PDGFRA, SOX17, FOXA2 and GATA4 (Fig. 2C). We co-stained for SOX2 or NANOG, since these epi markers are known to be lost in hypo precursors earlier than OCT4 (Blakeley et al., 2015; Chen et al., 2009; Niakan and Eggan, 2013; Roode et al., 2012) (Fig. 1,2B, Fig. S3E). PDGFRA appears in the membrane of all cells of the early ICM at D5, co-expressed with both SOX2 and NANOG (Fig. 3A-D). At early D6, emerging epi precursors downregulate PDGFRA and it becomes completely lost in the epi at late D6 while remaining strongly expressed in the hypo (Fig. 3A-D). This is the first indication of a common precursor in the ICM which gives rise to epi and hypo lineages. Distribution of SOX17 is heterogeneous at D5 and then widespread in all co-expressing ICM cells at early D6 (Fig. 1). SOX17 becomes specific to hypo at late D6 and it is still present at D7 (Fig. 3C). Neither FOXA2 nor GATA4 are expressed as early as D5 (Fig. 3). *FOXA2* expression is detectable earlier than *GATA4* at D6 based on scRNAseq analysis and image quantification (Fig. 3B,D-H). Moreover, FOXA2+GATA4-cells are apparent at D6. GATA4+ cells are exclusively found facing the cavity (Guo et al., 2021; Meistermann et al., 2021; Roode et al., 2012), suggesting its expression commences after physical epi:hypo, sorting, while FOXA2+ cells can be observed within the ICM core (Fig. 3B,C). FOXA2 and GATA4 differ with regard to co-expression with epi markers. High expressing FOXA2 cells have low levels of SOX2 and are negative for NANOG. In contrast, cells expressing GATA4 are able to co-express low levels of both SOX2 and NANOG (Fig. 3B-D,G,H). The difference in co-expression with NANOG suggests that despite its earlier appearance in the ICM cells, FOXA2 is a marker of a more specified hypo state. This could be an indication of ICM differentiation taking place asynchronously in a similar way to the mouse embryo (Saiz et al., 2016) or via an independent mechanism governing the specification of the hypoblast, such as exposure to the cavity or signalling molecules.

**Figure 3.**
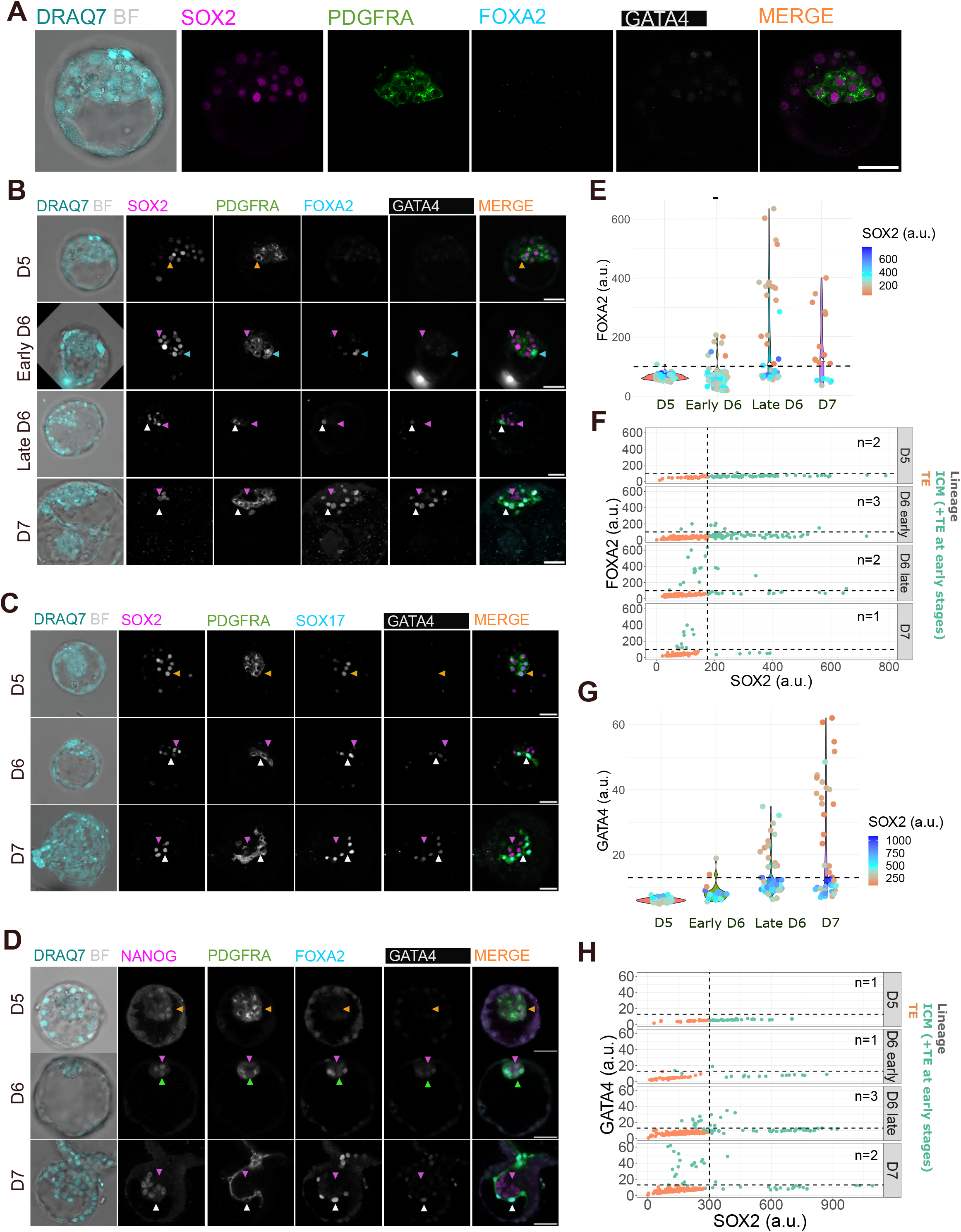
Validation of hypoblast marker order of appearance via protein characterization using immunostaining. **A**. Representative confocal image of a D5 human embryo showing PDGFRA (green) expression in all ICM cells, with subset labelled by nuclear SOX2 (magenta) expression as well. Neither FOXA2 nor GATA4 is expressed. Nucleus is visualized with DRAQ7 (turquoise). Scale bar = 50 µm. **B.C.D**. Representative confocal images of human embryos across blastocyst development D5 to D7 immunostained for epiblast markers SOX2 (A-C) or NANOG (D) in magenta and hypoblast markers PDGFRA (A-D) in green, GATA4 (A-D) in white, SOX17 (C) or FOXA2 (B,D) in cyan. Orange triangles indicate co-expressing cells, magenta triangles indicate epi precursors and epi cells. Cyan triangles indicate hypo precursors yet to express GATA4 (FOXA2+ GATA4-) while green triangles also indicate hypo precursors, positive for GATA4, negative for FOXA2 and still able to express epi markers such as NANOG. White triangles indicate hypo cells. Scale bar = 50 µm. **E**. Violin plot quantifying the nuclear expression of FOXA2 across different stages and its co-expression with nuclear SOX2 in the colour map. The dashed line represents the threshold calculated at D7 when hypo and epi have segregated into two lineages. **F**. Scatter plot showing quantification of nuclear intensity levels of SOX2 (epiblast marker) and FOXA2 (hypoblast marker). **G**. Violin plot quantifying the nuclear expression of GATA4 across different stages and its co-expression with nuclear SOX2 in the colour map. The dashed line represents the threshold calculated at D7 when hypo and epi have segregated into two lineages. **H**. Scatter plot showing quantification of nuclear intensity levels of SOX2 (epiblast marker) and GATA4 (hypoblast marker).

The presence of PDGFRA in the membrane of all ICM cells at early D5 (Fig. 3A-D) suggests that both epi and hypo populations arise from a common pool of ICM cells (Radley et al., 2022), rather than derivation of hypo cells from epi (Meistermann et al., 2021). Inclusion of SOX2 and NANOG in our characterization allowed us to determine which population between epi and hypo was specified first from the ICM. Already at D6, it is possible to see OCT4+SOX17-cells before OCT4-SOX17+ are present (Fig. 1A), implying the epi is specified first, as suggested previously (Boroviak et al., 2018). Furthermore, the protein characterization with both SOX2 and NANOG showed that SOX2+PDGFRA-as well as NANOG+PDGFRA-cells are present at D6, confirming that epi precursors appear earlier in development than the hypo counterpart (Fig. 3B-D). Low levels of SOX2 can be seen in hypo precursors until late D6 and NANOG is still expressed in PDGFRA+GATA4+ at D6, albeit not in cells already expressing high levels of FOXA2 (Fig. 3B-D).

To summarise, our data support the model in which epi and hypo populations arise from a common progenitor, the early ICM (Radley et al., 2022). While this conclusion was primarily based on existence of early ICM markers, such as LAMA4, that are downregulated in the differentiating ICM, here we investigated hypo marker expression at early stages to determine whether a co-expressing ICM population exists that subsequently resolves into separate lineages. The presence of PDGFRA in all ICM cells at D5 together with OCT4, SOX2 and NANOG indicates that both lineages originate in a population of cells co-expressing epi and hypo markers (Fig. 3,4). Our data also strengthen the hypothesis that epi is the first population to be specified, since NANOG+ only or SOX2+ only cells appear earlier than hypo+ only cells (Fig. 1,3,4), consistent with conclusions drawn from cell transfer experiments in preimplantation mouse embryos (Grabarek et al., 2012). Unfortunately, such definitive experiments are not possible using human embryos.

**Figure 4.**
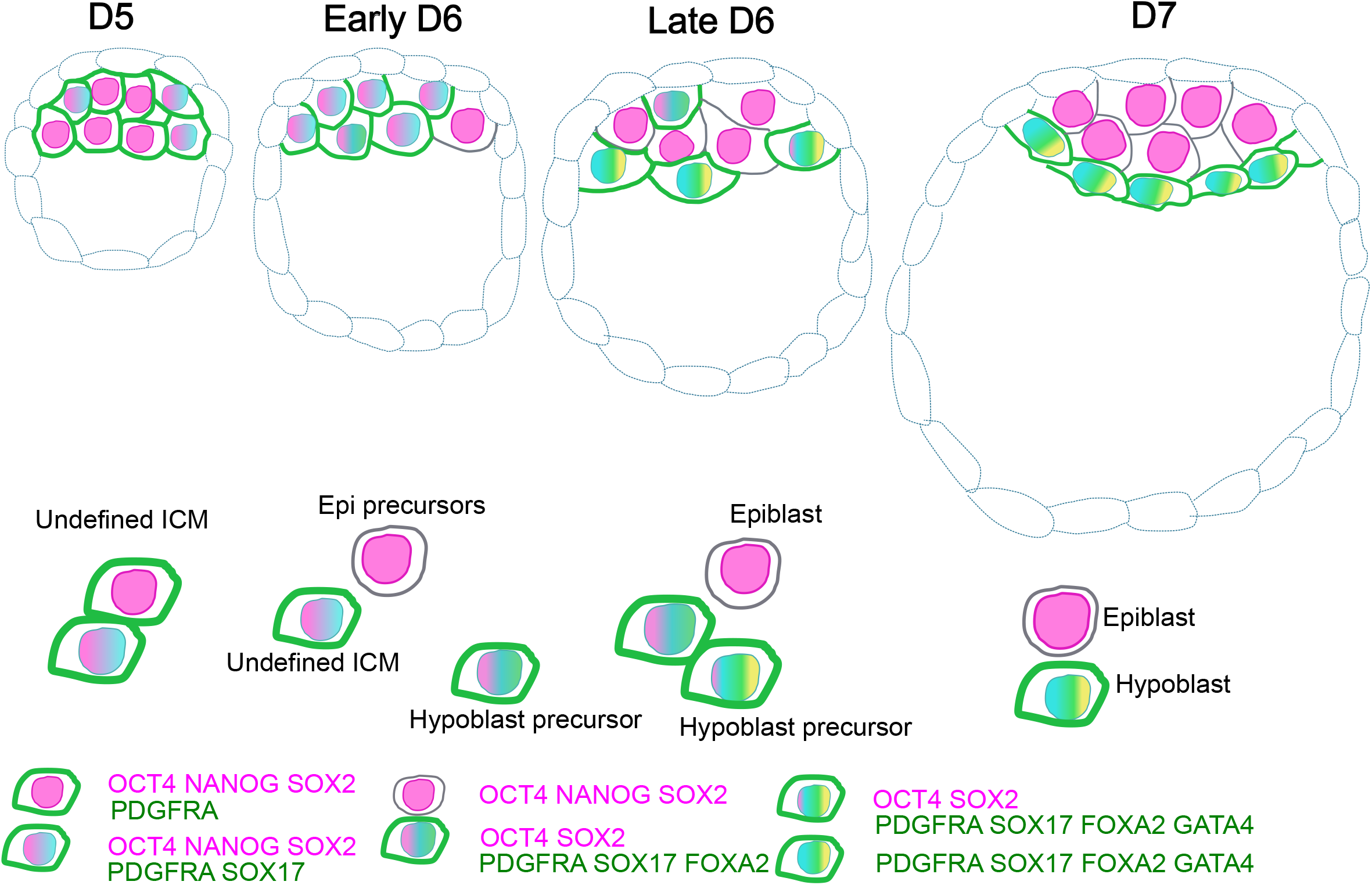
Schematic representation of the appearance of epiblast and hypoblast lineages in the human blastocyst based on the sequential “de novo” expression of hypoblast markers. At D5, the ICM comprises cells co-expressing the epi markers OCT4, NANOG and SOX2 together with the hypo marker PDGFRA in the cell membrane. At early D6, SOX17 expression begins in the nucleus of the undefined ICM population together with membrane PDGFRA, OCT4, SOX2 and NANOG. In parallel, some cells lose PDGFRA in the membrane and become epi precursors. At late D6, more epi cells have emerged within the ICM compartment. This stage is characterized by the appearance of hypoblast precursors: cells that express SOX17 and FOXA2 in the nucleus as well as PDGFRA in the membrane but have downregulated epi markers such as NANOG or SOX2. More advanced hypo cells can be found located next to the blastocoel expressing GATA4 in the nucleus, the last marker of this series to be expressed. At D7, both lineages are specified and sorted. The hypo cells are located next to the blastocoel and the epi compartment is situated between these and the polar TE. Epi cells express NANOG, OCT4 and SOX2 while hypo cells express PDGFRA, SOX17, FOXA2 and GATA4.

SOX2 can be maintained in human hypo cells until D6 but NANOG expression appears to be incompatible with FOXA2 expression arising in the embryo as early as D6 (Fig. 3) (Blakeley et al., 2015). The early specification of the epi could be related to epi factors already expressed in the early ICM, while the hypo population is strengthened with *de novo* expression of markers from D5 to D7. At D5 only PDGFRA and SOX17 are expressed, in contrast to the high levels of PDGFRA, SOX17, FOXA2 and GATA4 at D7 (Fig. 1, Fig. 3). As more antibodies for less well-known hypoblast markers become available, the process of lineage segregation in the developing human ICM can be further refined to uncover the optimal sequence for healthy blastocyst development.

## Materials and methods

### Human embryos

Supernumerary frozen human embryos were donated with informed consent by couples undergoing□*in*□*vitro*□fertility treatment. Use of human embryos in this research is approved by the Multi-Centre Research Ethics Committee, approval O4/MRE03/44, IRAS and licensed by the Human□Embryology□& Fertilization Authority of the United Kingdom, research license R0178.

Supernumerary frozen blastocysts (D5 and D6) were thawed and cultured in N2B27 medium (made in house) under mineral oil (Ovoil) in a humidified incubator at 37°C, 7% CO_2_ and 5% O_2_ until reaching the desired stage of development from E5 to E7. Embryonic stage was assessed based on thinning of the zona pellucida and blastocoele expansion (Fig. S1).

### Immunostaining and imaging of human embryos

The zona pellucida of D5 and D6 blastocysts was removed using acid Tyrode’s solution (Gibco) before fixation with 4% PFA in PBS for 15 min at room temperature. Embryos were rinsed in PBS containing 3 mg/ml polyvinylpyrrolidone (Sigma-Aldrich) (PBS/PVP), permeabilised using 0.25% Triton X-100 (Sigma-Aldrich) in PBS/PVP for 30 min and blocked in blocking buffer comprising PBS supplemented with 0.1% BSA, 0.01% Tween20 (Sigma-Aldrich) and 2% donkey serum for 2 h at room temperature. Primary and secondary antibodies were diluted in blocking buffer (see Table 1) Embryos were incubated in primary antibody solution overnight at 4°C and rinsed three times for 15 min in blocking buffer before incubation in secondary antibody solution for 1-2 h at room temperature in the dark. Embryos were rinsed in blocking buffer and imaged through a Poly-D-Lysin coated Mattek dish (P356-0-14) whilst submerged in blocking buffer. Embryos were imaged in a Leica Stellaris Confocal microscope and image analysis was performed using FIJI.

**Table 1.** Antibodies used for immunofluorescence.

| Antibody | Supplier | Cat no | Dilution |
| --- | --- | --- | --- |
| PDGFRα | Abcam | #ab203491<br>RRID:AB_ | 1:200 |
| SOX17 | R&D System | #AF1924<br>RRID:AB_355060 | 1:200 |
| GATA4 | EBioscience | #14-9980-82<br>RRID:AB_763541 | 1:200 |
| FOXA2/HNF-3β | R&D System | #AF2400<br>RRID:AB_2294104 | 1:200 |
| SOX2 | Santa Cruz | #sc-365823<br>RRID:AB_10842165 | 1:200 |
| OCT4? | Santa Cruz | #sc-5279<br>RRID:AB_628051 | 1:200 |
| DRAQ 7<sup>TM</sup> | Thermo Fisher<br>Scientific | #D15105 | 1:1000 |
| Donkey anti-goat 405 | Abcam | #ab175664<br>RRID:AB_2313502 | 1:500 |
| Alexa Fluor donkey<br>anti-rabbit 488 | Thermo Fisher<br>Scientific | #A32790<br>RRID:AB_2762833 | 1:500 |
| Alexa Fluor donkey<br>anti-mouse 555 | Thermo Fisher<br>Scientific | #A-31570<br>RRID:AB_2536180 | 1:500 |
| Alexa Fluor donkey<br>anti-rat 647 | Thermo Fisher<br>Scientific | #A78947<br>RRID:AB_2910635 | 1:500 |

### Image acquisition and quantification

Embryos were imaged in a Leica Stellaris 8 confocal microscope.

Confocal images were converted to 8-bit using FIJI. The Hoechst/nuclei channel was used for nuclei segmentation in 3D using ZeroCostDL4Mic StarDist via Google colaboratory (Schmidt et al., 2018; Weigert et al., 2018). For details on software training for embryo images, Kraunsoe et al., (in press).

The segmented image outcome was merged to the original image using FIJI. Integrated density, Mean fluorescence Intensity and volume per nuclei were measured using FIJI. Data analysis and data representation was performed using R. Scatter and violin plots were generated using the function “ggplot2” while histogram plots were created using “ggExtra”. No statistical analysis was performed due to the low number of embryos per stage per batch.

The expression threshold that separates the TE and ICM, and the ICM into epi, hypo and precursors was calculated based on the nuclear fluorescence intensity via generating histograms from the data at D7. At this late stage, epi and hypo markers are only expressed in the ICM, not TE; moreover, the ICMs are solely formed by epi and hypo with complete different marker expression meaning that plotting SOX17 and/or OCT4 generates at least two distinct normal distributions (negative and positive). The point in between the two normal distributions is considered the threshold and it is used in early stages to separate ICM from TE, and subsequently Epi and Hypo precursors from the undefined ICM population. Because OCT4 is expressed in the TE at D5, the ‘ICM’ population at D5 is misleadingly large in the scatter plot since it also contains the TE cells.

### Single cell RNA sequencing analysis

The single cell RNA sequencing analysis in this work was primarily built upon the high-resolution human pre-implantation embryo UMAP generated recently (Radley et al., 2022). A detailed workflow for reproducing the plots in this paper can be found at: https://github.com/aradley/Hypoblast_Activation_Paper.

In brief, we took the UMAP embedding defined by Radley et al., 2022 and focussed on the cells present in the bifurcation from the morula to the epi/hypo cell stages.

Pseudotime along the hypo branch was calculated using the Slingshot R packaged (Street et al., 2018).

Gene expression values were smoothed by taking the average gene expression for each cell and its 30 most similar cells according to the 3700 highly structured genes identified by Radley et al., 2022.

Logistic regression curves were fit to the smoothed expression profiles versus the hypo pseudotime. The hypo pseudotime value that corresponds to the half way point on the y-axis of the logistic curve indicates the tipping point between a gene being active Vs inactive.

## Author contributions

Conceptualization: J.N., E.C.S.; Methodology: E.C.S., A.R., J.N.; Software: A.R.; Formal analysis: A.R.; Investigation: E.C.S., A.R., J.N.; Resources: J.N.; Data curation: E.C.S.,J.N.; Writing - original draft: E.C.S.,A.R.,J.N.; Writing - review & editing: J.N., E.C.C., A.R.; Visualization: E.C.S., A.R.; Supervision: J.N.; Project administration: J.N.; Funding acquisition: J.N.

### Acknowledgement

We are grateful to Kenneth Jones, Ayaka Yanagida and Lawrence Bates for assistance with human embryo thawing. We are thankful to Peter Humphreys, Darran Clement Ann Wheeler and Mathew Pearson for help with microscopy, and Sophie Kraunsoe for assistance in image analysis.

## Funding

E.C.-S. was funded by the BBSRC grant number BB/T007044/2. A.R. was funded by a BBSRC PhD studentship (1943266) with co-funding from the Microsoft Research PhD scholarship program. JN was funded by the universities of Cambridge and Edinburgh.

## Competing interest

No competing interests declared.

## Supplementary figure legends

**Figure Supplementary 1. Human embryo staging from D5 to D7**. Representative bright field image of human embryos at D5, early D6, late D6 and D7. Differences in global size and blastocoel size were used to stage the embryos together with zona thickness (removed prior to image capture).

**Figure Supplementary 2. Additional imaging batches were examined to confirm the timing of hypoblast appearance in human embryos**. Batch 2: **A-C**; Batch 3: **D-F**; Batch 4: **G-I. A**,**D**,**G**. Scatter plots quantifying the nuclear intensity of OCT4 (x-axis) and SOX17 (y-axis) throughout blastocyst stages in the ICM (light green) and TE (orange) in individual cells. At early stages (D5 and early D6), light green also comprises TE cells due to OCT4 expression in the TE at the same level as ICM. Dashed lines represent the expression threshold calculated at D7. “n” refers to number of embryos analysed per stage. **B**,**E**,**H**. Scatter plots combining the quantification of OCT4 and SOX17 nuclear intensity per cell throughout all of the stages in blastocyst development (D5 to D7). Marginal density plots on the side and on top of the scatter plot show the data distribution that varies between discrete or heterogeneous populations. **C**,**F**,**I**. Violin plots showing the changes in SOX17 nuclear intensity across blastocyst stages and its co-expression with OCT4 per cell (colour-map). The dashed line represents the expression threshold for SOX17.

**Figure Supplementary 3. Marker expression across all cells of the human pre-implantation embryo, E3-E7. A, B**. Human pre-implantation embryo UMAP taken from Radley et al., 2022 where samples are labelled either by the cell type labels provided by Stirparo et al., 2018 (A) or manually labelled (B) based on the UMAP, cell type marker expression and the labels provided by Stirparo et al., 2018. **C-F**. Violin plots for different marker groups of interest, based of the groupings in B. **C**. Hypoblast markers that were presented in Figure 2, ordered from left to right in by their order of activation along the hypoblast pseudotime. **D**. Trophectoderm markers. **E**. Epiblast markers. **F**. Negative control with low expression throughout the human pre-implantation embryo.

**Figure Supplementary 4. Logistic regression of gene expression profiles along the hypoblast pseudotime**. We use the hypoblast pseudotime generated through the Slingshot software (Figure 2A) to fit a logistic curve to the expression profiles of genes of interest. A useful output of using a logistic curve is that the results provide a clear rationale for separating cells where a gene is considered inactive from cells where a gene is considered active. To fit a logistic curve, we first smooth the raw expression values (grey) of each cell by looking at their neighbourhood of k=30 cells and taking the average expression. We then use these smoothed expression values (blue) to fit a logistic curve, using the scipy.optimise.curve_fit function in Python. The resulting logistic curves are shown in red.

## Notes

### Competing Interest Statement

The authors have declared no competing interest.

## References

Artus, J., Piliszek, A., and Hadjantonakis, A.K. (2011). The primitive endoderm lineage of the mouse blastocyst: sequential transcription factor activation and regulation of differentiation by Sox17. Dev Biol 350, 393–404.

Blakeley, P., Fogarty, N.M., Del Valle, I., Wamaitha, S.E., Hu, T.X., Elder, K., Snell, P., Christie, L., Robson, P., and Niakan, K.K. (2015). Defining the three cell lineages of the human blastocyst by single-cell RNA-seq. Development 142, 3151–3165.

Boroviak, T., Stirparo, G.G., Dietmann, S., Hernando-Herraez, I., Mohammed, H., Reik, W., Smith, A., Sasaki, E., Nichols, J., and Bertone, P. (2018). Single cell transcriptome analysis of human, marmoset and mouse embryos reveals common and divergent features of preimplantation development. Development 145.

Chazaud, C., Yamanaka, Y., Pawson, T., and Rossant, J. (2006). Early lineage segregation between epiblast and primitive endoderm in mouse blastocysts through the Grb2-MAPK pathway. Dev Cell 10, 615–624.

Chen, A.E., Egli, D., Niakan, K., Deng, J., Akutsu, H., Yamaki, M., Cowan, C., Fitz-Gerald, C., Zhang, K., Melton, D.A., et al. (2009). Optimal timing of inner cell mass isolation increases the efficiency of human embryonic stem cell derivation and allows generation of sibling cell lines. Cell Stem Cell 4, 103–106.

Gerri, C., McCarthy, A., Alanis-Lobato, G., Demtschenko, A., Bruneau, A., Loubersac, S., Fogarty, N.M.E., Hampshire, D., Elder, K., Snell, P., et al. (2020). Initiation of a conserved trophectoderm program in human, cow and mouse embryos. Nature 587, 443–447.

Grabarek, J.B., Zyzynska, K., Saiz, N., Piliszek, A., Frankenberg, S., Nichols, J., Hadjantonakis, A.K., and Plusa, B. (2012). Differential plasticity of epiblast and primitive endoderm precursors within the ICM of the early mouse embryo. Development 139, 129–139.

Guo, G., Stirparo, G.G., Strawbridge, S.E., Spindlow, D., Yang, J., Clarke, J., Dattani, A., Yanagida, A., Li, M.A., Myers, S., et al. (2021). Human naive epiblast cells possess unrestricted lineage potential. Cell Stem Cell 28, 1040–1056 e1046.

Hertig, A.T., Rock, J., and Adams, E.C. (1956). A description of 34 human ova within the first 17 days of development. Am J Anat 98, 435–493.

Kuijk, E.W., van Tol, L.T., Van de Velde, H., Wubbolts, R., Welling, M., Geijsen, N., and Roelen, B.A. (2012). The roles of FGF and MAP kinase signaling in the segregation of the epiblast and hypoblast cell lineages in bovine and human embryos. Development 139, 871–882.

Meistermann, D., Bruneau, A., Loubersac, S., Reignier, A., Firmin, J., Francois-Campion, V., Kilens, S., Lelievre, Y., Lammers, J., Feyeux, M., et al. (2021). Integrated pseudotime analysis of human pre-implantation embryo single-cell transcriptomes reveals the dynamics of lineage specification. Cell Stem Cell.

Mole, M.A., Weberling, A., Fassler, R., Campbell, A., Fishel, S., and Zernicka-Goetz, M. (2021). Integrin beta1 coordinates survival and morphogenesis of the embryonic lineage upon implantation and pluripotency transition. Cell Rep 34, 108834.

Niakan, K.K., and Eggan, K. (2013). Analysis of human embryos from zygote to blastocyst reveals distinct gene expression patterns relative to the mouse. Dev Biol 375, 54–64.

Nichols, J., Silva, J., Roode, M., and Smith, A. (2009). Suppression of Erk signalling promotes ground state pluripotency in the mouse embryo. Development 136, 3215–3222.

Nowotschin, S., Setty, M., Kuo, Y.Y., Liu, V., Garg, V., Sharma, R., Simon, C.S., Saiz, N., Gardner, R., Boutet, S.C., et al. (2019). The emergent landscape of the mouse gut endoderm at single-cell resolution. Nature 569, 361–367.

Petropoulos, S., Edsgard, D., Reinius, B., Deng, Q., Panula, S.P., Codeluppi, S., Plaza Reyes, A., Linnarsson, S., Sandberg, R., and Lanner, F. (2016). Single-Cell RNA-Seq Reveals Lineage and X Chromosome Dynamics in Human Preimplantation Embryos. Cell.

Plusa, B., Piliszek, A., Frankenberg, S., Artus, J., and Hadjantonakis, A.K. (2008). Distinct sequential cell behaviours direct primitive endoderm formation in the mouse blastocyst. Development 135, 3081–3091.

Radley, A., Corujo-Simon, E., Nichols, J., Smith, A., and Dunn, S.J. (2022). Entropy sorting of single-cell RNA sequencing data reveals the inner cell mass in the human pre-implantation embryo. Stem Cell Reports.

Roode, M., Blair, K., Snell, P., Elder, K., Marchant, S., Smith, A., and Nichols, J. (2012). Human hypoblast formation is not dependent on FGF signalling. Dev Biol 361, 358–363.

Saiz, N., Mora-Bitria, L., Rahman, S., George, H., Herder, J.P., Garcia-Ojalvo, J., and Hadjantonakis, A.K. (2020). Growth-factor-mediated coupling between lineage size and cell fate choice underlies robustness of mammalian development. Elife 9.

Saiz, N., Williams, K.M., Seshan, V.E., and Hadjantonakis, A.K. (2016). Asynchronous fate decisions by single cells collectively ensure consistent lineage composition in the mouse blastocyst. Nat Commun 7, 13463.

Schmidt, U., Weigert, M., Broaddus, C., and Myers, G. (2018). Cell Detection with Star-Convex Polygons (Cham: Springer International Publishing).

Stirparo, G.G., Boroviak, T., Guo, G., Nichols, J., Smith, A., and Bertone, P. (2018). Integrated analysis of single-cell embryo data yields a unified transcriptome signature for the human pre-implantation epiblast. Development 145.

Street, K., Risso, D., Fletcher, R.B., Das, D., Ngai, J., Yosef, N., Purdom, E., and Dudoit, S. (2018). Slingshot: cell lineage and pseudotime inference for single-cell transcriptomics. BMC Genomics 19, 477.

Weigert, M., Schmidt, U., Boothe, T., Muller, A., Dibrov, A., Jain, A., Wilhelm, B., Schmidt, D., Broaddus, C., Culley, S., et al. (2018). Content-aware image restoration: pushing the limits of fluorescence microscopy. Nat Methods 15, 1090–1097.

Yan, L., Yang, M., Guo, H., Yang, L., Wu, J., Li, R., Liu, P., Lian, Y., Zheng, X., Yan, J., et al. (2013). Single-cell RNA-Seq profiling of human preimplantation embryos and embryonic stem cells. Nat Struct Mol Biol 20, 1131–1139.

Zhu, M., and Zernicka-Goetz, M. (2020). Principles of Self-Organization of the Mammalian Embryo. Cell 183, 1467–1478.

